# Connecting gene expression and cytokinin signaling during meristem determinacy transition in the rice panicle

**DOI:** 10.64898/2026.04.17.719148

**Authors:** James Tregear, Carole Gauron, Virginie Vaissayre, Frances N Borja, Julien Serret, Caroline Michaud, Marie-Christine Combes, Guillaume Cazals, Daphné Autran, Stefan Jouannic, Hélène Adam

## Abstract

Structural complexity in the rice inflorescence (panicle) is determined by the activity of indeterminate meristems, which allow sequential branching events to occur but later acquire a determinate character that precludes the initiation of new growth axes. To better understand the underlying regulatory processes, we combined a detailed time-course of panicle development with tissue-specific sampling of meristems before and after the determinacy switch, monitoring global gene expression programs and hormone accumulation. This allowed the delineation of three dynamic transcriptional modules and the inference of gene regulatory networks highlighting hormone regulatory hubs associated with the indeterminate-to-determinate meristem transition. Our data confirm the importance of known regulators of inflorescence development and identify novel actors, notably genes encoding transcription factors, that were not previously described in the regulation of flowering. The combined biochemical and transcriptomic data support a model in which cytokinin signalling is particularly active during the proliferative branching phase, during which meristem maintenance is promoted while a feedback mechanism is also triggered, ultimately leading to the acquisition of determinate meristem fate later in development.

**Highlight:** Investigating the molecular nature of rice inflorescence meristem determinacy, we studied temporal and spatial gene expression alongside hormone accumulation, identifying novel regulatory interactions and a key role for cytokinins.

## 1 Introduction

The transition of plant meristem identity from an indeterminate to a determinate state is a crucial process in development, governing growth patterns and morphology. Following floral induction, inflorescence development begins with the transition of a vegetative shoot apical meristem (SAM) into the inflorescence rachis meristem (RM). In the case of the rice panicle, a compound inflorescence (Fig. 1), the RM produces axillary meristems (AMs) that develop as primary branch meristems (PbMs). The latter then generate their own AMs, which often display indeterminate meristem (IM) identity, i.e., act as branch meristems (BMs) to allow secondary and sometimes tertiary branching. IM identity is ultimately lost, and determinate meristem (DM) identity is acquired, culminating in spikelet meristem (SpM) development. In the later stages of panicle development, all IMs transition gradually into a DM state from the top to the base of the panicle and from the terminal meristem to the basal part of the branches, resulting in the appearance of SpMs and then florets (Ikeda et al., 2004; Itoh et al., 2005; Zhang and Yuan, 2014). The timing of the shift from an indeterminate to a determinate state is crucial, directly affecting the panicle’s shape, density, and resource capture efficiency for seed development (Kyozuka et al., 2014; Wang et al., 2021). Understanding this transition is essential to explaining how plants manage growth and reproduction in response to internal and external cues.

**Fig. 1:**
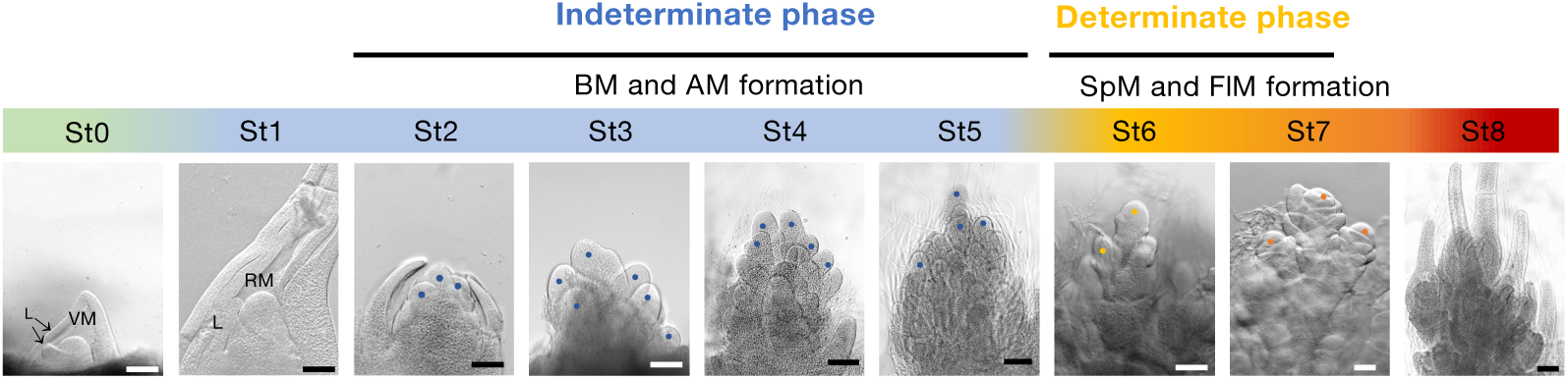
Early stages of rice panicle development analyzed for transcriptomics and hormone profiling. Sampling of immature panicle at nine developmental stages; St0 – vegetative meristem (VM); St1 - rachis meristem (RM); St2 - primary branch meristem (PbM); St3 – early primary branch elongation; St4 – primary branch elongation with axillary meristems (AMs); St5 - primary branch elongation with secondary branch meristem (SBM); St6 – Spikelet meristem (SpM); St7 – Floret meristem (FlM); St8 – Floral organ (FlO) initiation. Scale bars: 100 μm. Coloured dots indicate meristem types as follows: blue, terminal PbM; yellow, SpM; orange, FlM.

Previous studies on rice have highlighted the pivotal role of specific transcription factor (TF) gene families during panicle differentiation (Tanaka et al., 2023). Numerous *APETALA2*-*like* (*AP2-like*) gene functions are associated with the branching stage of inflorescence development, whereas those of MADS box genes are preponderantly associated with the determinate phase (Furutani et al., 2006; Harrop et al., 2016; Harrop et al., 2019; Zong et al., 2022). Similarly, the rice TF gene *TAWAWA1* (*TAW1*; Yoshida et al., 2013) regulates meristem activity during panicle development by promoting the indeterminate activity of the AMs, thereby delaying the loss of indeterminacy that characterises SpMs. TAW1 belongs to the ALOG TF family, containing a highly conserved domain of major importance in plant growth and development (Naramoto et al., 2020; Lyer and Aravind, 2012). Another key player is the SQUAMOSA PROMOTER-BINDING PROTEIN-LIKE (*SPL*) gene *OsSPL14*, also known as *IDEAL PLANT ARCHITECTURE1* (*IPA1*)/ *WEALTHY FARMER’S PANICLE* (*WFP*), encoding a TF that promotes panicle branching, leading to a higher grain yield (Jiao et al., 2010; Miura et al., 2010). The aforementioned activities are opposed by several TF-encoding genes that promote the transition from indeterminate to determinate meristem identity and the subsequent transition from the spikelet to the floret meristem. Examples in this group include the MADS-box gene *LEAFY HULL STERILE* (*LHS1/OsMADS1*), along with the *AP2-like* genes *FRIZZY PANICLE* (*FZP*) and *MULTI-FLORET SPIKELET1* (*MFS1)* (Komatsu et al., 2003; Agrawal et al., 2005; Khanday et al., 2013; Ren et al., 2020).

Hormones such as auxins and cytokinins play a significant role in the regulation of meristem identity and transition, their actions being closely entwined with those of many of the abovementioned TFs (Deveshwar et al., 2020). From a broad perspective, auxins have been extensively implicated in AM initiation, while cytokinins play a key role in the regulation of meristem size and activity, which in turn affects branching, notably in the case of the cereal panicle. *Gn1a*, a major panicle branching QTL, encodes the cytokinin oxidase/dehydrogenase OsCKX2, which degrades cytokinins. Reduced expression of *Gn1a/OsCKX2* leads to increased cytokinin accumulation in the RM, promoting higher branch numbers and an enhanced grain yield (Ashikari et al., 2005). Conversely, a loss-of-function mutation in the *LONELY GUY* (*OsLOG*) gene, which encodes an enzyme responsible for cytokinin activation, results in premature termination of shoot and inflorescence meristematic activities, causing the formation of small, underdeveloped panicles (Kurakawa et al., 2007). Further studies have shown that *OsSPL14* contributes to the regulation of panicle architecture through cytokinin signalling. The OsSPL14 protein binds directly to the promoter of *LOG*, thus promoting cytokinin biosynthesis (Lu et al., 2013), while also regulating *DENSE AND ERECT PANICLE 1* (*DEP1*), a key determinant of grain number per panicle (Yan et al., 2007; Huang et al., 2009; Li et al., 2025). *DEP1* activation is associated with downregulation of *OsCKX2*, further increasing cytokinin levels in meristems (Huang et al., 2009; Lu et al., 2013; Zhang et al., 2025). Cytokinin levels in the inflorescence are also regulated by other factors. For example, genes such as *DROUGHT AND SALT TOLERANCE* (*DST*), *LARGER PANICLE* (*LP*), and *VIN3-LIKE 2* (*OsVIL2*) influence panicle architecture and cytokinin homeostasis by modulating *Gn1a/OsCKX2* expression (Li et al., 2013; Li et al., 2011; Yang et al., 2019). More recently, *FZP* was shown to function upstream of the DST–OsCKX2 module, maintaining cytokinin homeostasis in the inflorescence meristem and regulating spikelet development (Wang et al., 2023). A link between cytokinins and inflorescence development was also demonstrated by Hirose et al. (2007) through transgenic overexpression of the rice cytokinin-inducible type A response regulator OsRR6, which resulted in an altered panicle morphology. Auxin also plays a central role in panicle patterning by being essential for the initiation and maintenance of axillary meristems. Panicle branching can be disrupted by perturbations in the auxin pathway, particularly through the alteration of genes involved in local auxin biosynthesis, transport, signalling, distribution or response (Zhang and Yuan 2014; Deveshwar et al., 2020). One such gene, *NAL1* (*NARROW LEAF1*), pleiotropically affects both plant and panicle architecture, as well as grain yield, via regulation of polar auxin transport (Qi et al., 2008). Similarly, Liu et al. (2021) demonstrated the critical role of the rice auxin transporters encoded by *OsPIN1c* and *OsPIN1d* during panicle development. Loss of function of these genes results in a “pin-shaped” panicle phenotype, accompanied by widespread deregulation of key regulators involved in reproductive development. The importance of spatial auxin and cytokinin distribution patterns in rice panicle development has also been highlighted using reporter gene constructs (Yang et al., 2017; Sato et al., 2025).

Despite extensive research on the rice panicle, the exact mechanisms by which hormonal crosstalk and TF regulation govern the transition between the IM and DM states remain incompletely understood. One approach that has proven useful is the physical dissection of component tissues to identify genes with specific spatial expression patterns. Previous studies using laser microdissection combined with RNA sequencing (RNA-seq) and single-cell analysis revealed gene expression profiles and clusters associated with early panicle development (Harrop et al., 2016; Zong et al., 2022). In the present study, we characterised temporal and tissue-specific molecular factors using a three-pronged approach: firstly, via a single panicle developmental time-course transcriptomic dataset; secondly, through panicle dissection to compare gene expression between the IM and DM states; and thirdly, by biochemical profiling of auxin/cytokinin accumulation in IM- and DM-phase panicles.

## 2 Materials and Methods

### 2.1-Plant materials and growth conditions

*O. sativa* spp. *japonica* cv. Nipponbare plants were grown in a greenhouse for 7 weeks in long-day conditions (14h/10h day/night cycle at 28/26°C and 60% humidity). Plants were then transferred into short-day conditions (10h/14h day/night cycle) for flowering induction.

### 2.2-Sampling, RNA extraction, and sequencing

For single panicle analysis, immature panicles were harvested from plants at nine defined developmental stages, namely: the vegetative meristem (VM) stage (St-0); the rachis meristem (RM) stage (St-1); four different indeterminate branch meristem (BM) stages (St-2 to St-5) and three different determinate branch meristem (DM) stages characterised by spikelet meristem (SpM), floret and floral organ formation (St-6, St-7 and St-8 respectively) (Fig. 1). Samples were manually dissected under a stereomicroscope (Leica S8APO) for imaging and stage determination (Supplementary Fig. S1). Each dissected panicle was photographed (Leica DFC295) and its developmental stage noted (St-0 to St-8), after which it was transferred to a Safe-Lock 2 mL tube (Eppendorf) containing a single 5-mm glass bead (Sigma–Aldrich). The tube was then transferred to liquid nitrogen and stored at −80 °C prior to use in RNA extraction. For RNA-seq analysis, total RNAs were extracted from each individual panicle sample using a RNeasy Micro kit (QIAGEN). Quantification and quality checks of the extracted RNAs were carried out using an Agilent Bioanalyzer to ensure that samples were undegraded (RNA integrity number ≥8). RNA-seq analysis was performed on three biological replicates per panicle stage. Synthesis of cDNA libraries was carried out from 100ng samples using the TruSeq Illumina Strand Specific protocol, the size of the selected indexed fragments being from 300 to 750 bp. Illumina (HiSeq 3000) sequencing was carried out by Novogene (Cambridge, UK) to obtain 150-base-pair paired-end reads.

For single meristem analysis, individual IM and DM samples were obtained by excising from the apex of the distal primary branch of a panicle at either the BM or SpM phase of development. Samples were placed individually into 0.5 mL tubes containing the extraction buffer. RNA was extracted using the ARCTURUS PicoPure RNA isolation 600 Pico Kit (Agilent, Santa Clara, CA, USA) with additional grinding of tissues using BEAD (Glasperlen 1.7-2.1mm, Roth). RNA yields were evaluated using TapeStation and the High Sensitivity RNA ScreenTape (Agilent). RNA-seq analysis was performed on three biological replicates per meristem type. One or two nanograms of RNA were used for cDNA library preparation after 11 amplification cycles using a NEBNEXT Single/low input RNA library kit (New England Biolabs). Library quality was assessed with a Bioanalyzer (Agilent) using the High Sensitivity DNA assay protocol. DNA sequencing was performed by Novogene (Cambridge, UK) using the Illumina NovaSeq platform, generating 150 nt paired-end reads.

### 2.3-Sequence data processing and gene expression analysis

The quality of each sequenced library was checked by analysing the corresponding reads obtained using FastQC (Andrews, 2010). Reads were trimmed using CutAdapt (Martin, 2011) according to the following criteria: the first 12 bases of all reads were eliminated, contaminating adapter sequences were removed; low-quality bases (Q<30) were trimmed out from both the 5- and 3-ends of reads; and reads less than 35 nucleotides were discarded. Cleaned reads were mapped against the *O. sativa* ssp. *japonica* cv. Nipponbare IRSGP1 reference genome (RAP-DB annotation database https://rapdb.dna.affrc.go.jp, version of 17/01/2024) using STAR with a mismatch maximum per read of 3% and the quantMode option GeneCounts to assign mapped reads to annotated loci (Dobin et al., 2013; Kawahara et al., 2013; Sakai et al., 2013). Count files from the individual samples were merged to obtain a count table for subsequent analyses. Single panicle and single meristem datasets were processed separately.

#### Gene expression analysis

Gene expression analyses were performed using the edgeR package integrated within the DIANE (Dashboard for the Inference and Analysis of Networks from Expression data) Shiny platform (Cassan et al., 2021) using TMM (Trimmed Means of the M-values) as the normalization tool and a cut-off filtering for the minimal gene count sum across the 3 replicate libraries of 30. Differentially expressed genes (DEGs) were identified using the criteria log_2_Fold Change (log_2_FC) ≥1 or ≤−1 and False Discovery Rate (FDR) of either <0.01 for the single panicle dataset, or <0.05 for the single meristem dataset.

#### Co-expression analysis

To analyse the dynamics of gene expression across branching stages, co-expression analysis was performed on the identified set of DEGs using a multifactorial model. Clustering was carried out using the coseq R package (Rau and Maugis-Rabusseau, 2018) implemented within the DiCoExpress R-based workspace (Lambert et al., 2020). Normalized expression data were transformed using an arcsin transformation prior to clustering. Clustering analyses were performed on the 1573 DEGs, corresponding to the union of DEGs from the successive contrasts St2-St3, St3-St4, St4-St5, St4-St6, St5-St6, and St6-St7. A Gaussian mixture model was fitted while testing a range of 3 to 20 clusters with a fixed random seed of 100. The optimal number of clusters was determined using the Integrated Completed Likelihood (ICL) criterion, and genes were grouped according to their expression profiles across all samples.

#### Functional enrichment analyses

For DEGs and genes in co-expressed clusters, gene ontology (GO term) analyses were performed using the R scripts and annotation file described by Li (2022), with an adjusted *p*-value threshold of 0.05 (Ashburner et al., 2020; The Gene Ontology Consortium, 2023; Kanehisa et al., 2020, 2025). Redundant GO terms were filtered using a dispensability cut-off (<0.05) (Supek et al., 2011). Transcription factor- and hormone-related gene enrichment was assessed using annotations from the Plant Transcription Factor Database PlnTFDB v5.0 (Tian et al., 2020; Jin et al., 2017) and from a curated gene set relating to hormone function (Zong et al., 2022), respectively, *via* a hypergeometric test with Benjamini–Hochberg correction (*p*-value < 0.05). Heatmaps were generated using the R package Pheatmap (Version 1.0.12).

#### Gene regulatory network inference

To focus on the regulation of meristem determinacy during panicle development, transcriptomic data from St2 to St7 were used as the basis for gene regulatory network (GRN) inference. A gene set consisting of DEGs identified between St4–St6 and St5–St6 in the single panicle dataset plus IM vs DM DEGs (|log2FC| > 2, FDR < 0.01) was initially assembled, retaining only DEGs that belonged to defined co-expression clusters. This 427 gene set was enriched with 10 additional expert-curated genes previously described as essential for panicle development, serving as biological anchors for the GRN analysis. Transcription factor- and hormone-related genes were defined as candidate regulators. GRN inference was carried out using the DIANE pipeline, which implements a regression tree–based random forest approach to reconstruct putative regulatory relationships (based on 1000 trees). Edge selection was performed using a network density cutoff of 0.03, and only interactions with an FDR < 0.05 were retained for downstream analyses. The final predicted network was imported into Cytoscape v3.8.0 (Shannon et al., 2003) to generate a high-quality graphical representation of the GRN. Node shapes, colours, and sizes were used to distinguish TFs from non-TFs and to indicate up- or down-regulation between the IM and DM phases. Network nodes were further partitioned into communities using the Louvain method for community detection (Blondel et al., 2008).

### 2.4-Auxin and cytokinin profiling using UPLC-MS/MS

Phytohormones were identified and quantified by ultraperformance liquid chromatography coupled with tandem mass spectrometry (UPLC-MS/MS) using MS transitions determined by means of pure standards. Indole-3-acetic acid (IAA), IAA-aspartic acid (IAA-Asp), IAA-alanine (IAA-Ala), IAA-glutamic acid (IAA-Glu), IAA-Leucine (IAA-Leu), and *cis-*/*trans*-Zeatin (Z-mix), were purchased from OlChemim Ltd. (Olomouc, Czech Republic), whereas *trans*-Zeatin-O-glucoside (tZOG), *trans*-Zeatin Riboside (tZR), isopentenyladenine 2 (2iP) and Isopentenyladenosine (iPA) were obtained from Santa Cruz biotechnology, Inc. (Dallas, Texas, USA). Stable isotope-labelled compounds [^2^H_5_]tZ, [^2^H_5_]tZR, [^2^H_6_]-iPA were purchased from OlChemim Ltd. and [^2^H_5_]-IAA from Cayman Chemicals (Ann Arbor, Michigan, USA). Internal standards were prepared as described by Vaissayre et al. (2025). Per replicate, four whole developing panicles at the BM or SpM phase were ground. For phytohormone extraction, 1 mL of methyl tert-butyl ether MTBE/Methanol (3:1, v/v) containing 50 ng of internal standards (Supplementary Table S1) was added to 25 mg of pre-weighted frozen sample powder in a 2 mL screw-cap tube. Samples were vortexed for 30 s, ultrasonicated for 15 min at 4°C and shaken on a rotary wheel for 30 min. Subsequently, 500 µL of 0.1% HCl was added, followed by 30 s of vortexing and 30 min of stirring at 4°C. After centrifugation at 10000 x g for 10 min at 4°C, 650 µL of supernatant was transferred to a 1.5 mL vial. The eluate was evaporated under a gentle stream of nitrogen and the pellet was reconstituted in 100 µL Methanol:H_2_0 (1:1, v/v) for direct analysis by UPLC-MS. Preparation of internal standard solution, instrument setup, data acquisition, and analysis were performed according to the protocol described in Vaissayre et al. (2025). The obtained results were statistically analysed with a *t*-test between the 2 stages and represented using the R package ggplot2 (version 3.5.1).

## 3 Results

### 3.1-Gene expression dynamics during early panicle development

We performed RNA-seq analysis on individual panicles harvested from *O. sativa* cv. Nipponbare plants at nine defined stages (St0-tSt8; Fig. 1 and Supplementary Fig. S1), spanning the complete trajectory from the vegetative meristem (VM) stage, through floral transition, into reproductive development, capturing the sequential establishment of branch meristems (BM) and their subsequent transition into determinate spikelet meristems (SpM). Our pipeline allowed the detection of transcripts of 26881 genes (Supplementary Tables S2, S3 and S4). Principal component analysis (PCA) performed on the corresponding expression data (Fig. 2A) confirmed distinct groupings of biological samples according to their developmental stage, although samples stage St4 and stage St5 exhibited partial overlap. The rachis meristem stage (RM; stage St1) was excluded from subsequent expression analyses, due to a higher expression variability being observed. Stage-specific differential expression was also visualised by pairwise comparisons of successive developmental stages (Fig. 2B, Supplementary Table S5), with respect to numbers of up- and down-regulated DEGs. Histograms of corresponding *p*-values obtained from differential expression analysis using an FDR threshold of 0.01 (Supplementary Fig. S2) confirmed robust statistical quality across all contrasts examined. Our analysis revealed that only a small number of DEGs (29 in total) distinguished stages St4 and St5, compared to 154 and 505 DEGs marking the transitions from stages St3 to St4 and from stages St5 to St6, respectively. Taken together, these results validated the sampling approach used to distinguish successive phases of panicle development, while revealing that the molecular distinction between stages St4 and St5, in terms of gene expression, is less marked compared to the other developmental transitions evaluated. The largest number of DEGs (533 in total) was observed in the comparison between stages St0 and St2, marking the crucial transition from vegetative to reproductive development. A similar number of DEGs (505 in total) characterized the transition from stage St5 to stage St6, during the shift from indeterminate to determinate meristem identity. GO enrichment analysis provided further information on the molecular nature of each developmental transition, in terms of biological process (BP) and molecular function (MF) (Supplementary Fig. S3A; Supplementary Table S6). The stage transition displaying the largest number of significantly affected biological process categories (7 out of 9) was that from stage St5 to stage St6. A high overall prevalence of TF genes (6.9% to 20%) was observed in the seven DEG sets (Fig. 2B). Several TF families, such as the LOB-LBD and CCAAT classes, showed a marked stage-specificity in their differential expression, consistent with their known importance in early flowering events (Xu et al., 2016). Distinct TF family enrichments were also observed in the St5–St6 transition DEG set, which exhibited significant overrepresentation of several TF families including the AP2-EREBP, MADS-box, LFY, ALOG, bHLH, HB and SPL classes (Supplementary Fig. S3B, Supplementary Table S7).

**Fig. 2:**
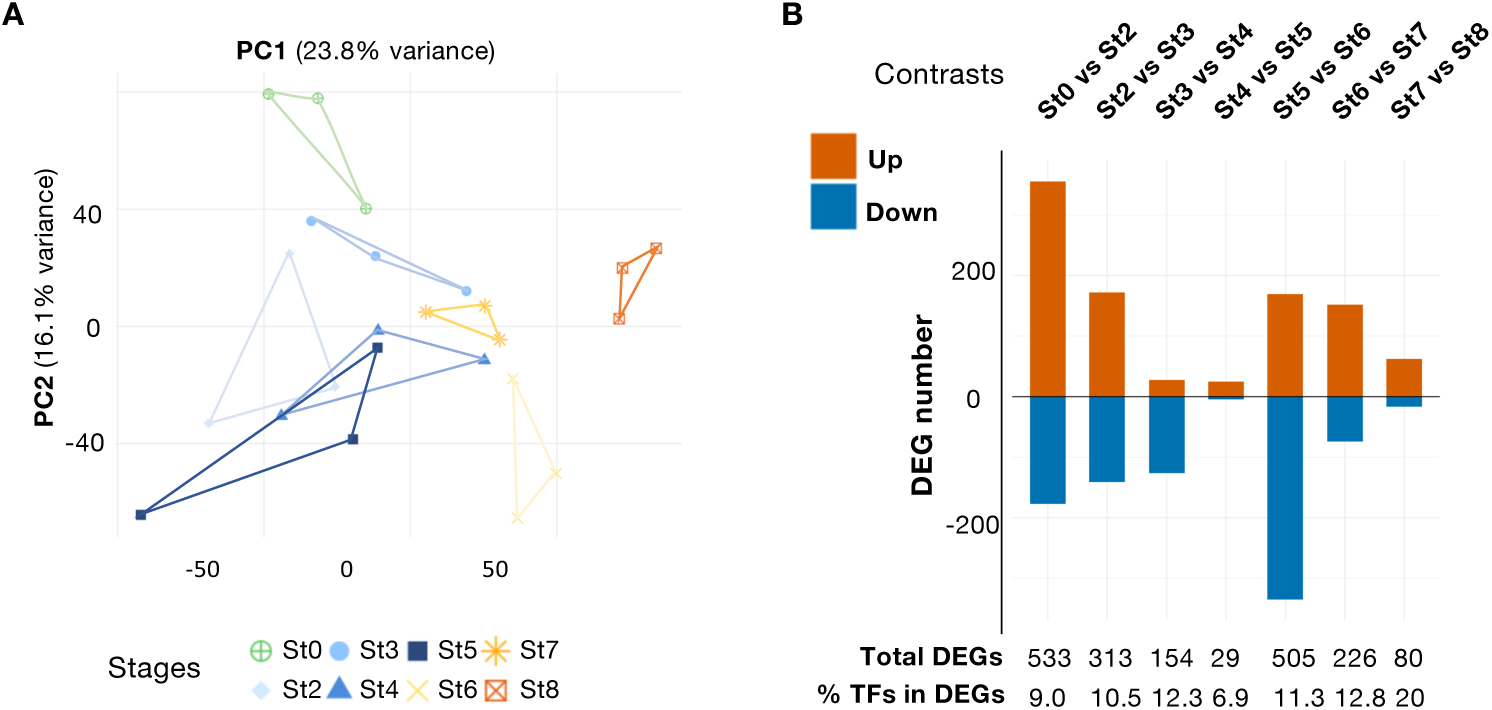
Transcriptomic analysis of early rice inflorescence development. **A.** Principal-component analysis (PCA) of gene expression data. Panicle developmental stages are distinguished by different colours and symbols as indicated. Three biological replicates were analyzed for each stage. **B.** Distribution of differentially expressed genes (DEGs) with respect to successive stage transitions in panicle differentiation. DEGs were identified using a false discovery rate (FDR) threshold of < 0.01 and an absolute |log₂ fold change| ≥ 1. Sequential pairwise comparisons (St0–St2, St2–St3, St3–St4, St4–St5, St5–St6, St6–St7, St7–St8) are presented. For each comparison, the number of upregulated (red) and downregulated (blue) genes, and the percentage proportion of DEGs encoding transcription factors (TFs) are shown.

We also assessed the presence of hormone-related genes in the DEG sets (Supplementary Fig. S3B, Supplementary Table S7) to provide information on their likely involvement in early panicle development. Our analysis indicated significant enrichment of cytokinin-related DEGs at the first three transitions (St0-2, St2-St3, and St3-St4), of auxin-related DEGs at two transitions (St0-St2 and St5-St6), and of brassinosteroid-related DEGs at the St5-St6 transition. Collectively, these data suggest that coordinated transcription factor activities and hormonal signaling pathways are major drivers of rice panicle differentiation and meristematic activity.

### 3.2-Co-expression analysis reveals three major transcriptional clusters characterising panicle differentiation

To identify gene groups associated with panicle differentiation, co-expression clustering was performed using all DEGs identified across the successive developmental stages from St2 to St7. Our analysis allowed the identification of three gene clusters displaying distinct expression dynamics during panicle development (Fig. 3A, Supplementary Fig. S4A). The ICL (Integrated Completed Likelihood) curve displayed a clear minimum, indicating a well-supported clustering structure (Supplementary Fig. S4B and S4C; cluster compositions shown in Supplementary Table S8). The three gene clusters displayed contrasting temporal profiles, but shared the common feature of a significant expression change between stages St5 and St6, i.e., within the developmental window during which indeterminate AMs progressively differentiate into spikelet meristems (Fig 3A). Enrichment analyses were conducted for each gene cluster with respect to GO categories plus TF- and hormone-related terms (Fig. 3B and 3C, Supplementary Tables S9 and S10). Cluster 1 (Cl1) consisted of 207 genes displaying a rising expression trend that peaked at stage St6, with a later increase at stage St8 (Fig. 3A). In contrast, Cl2 (362 genes) was characterized by an overall downward trend in gene expression, displaying a peak at stage St5 and a sharp decrease between stages St5 and St6, with a sub-peak at stage St7. Cl3 (757) genes displayed a flatter expression profile than the other two clusters, with an expression maximum at stage St3 and a slightly lower level at stage St8 relative to stage St0 (Fig. 3A). Interestingly, from stage St4 onwards, Cl2 and Cl3 displayed the same qualitative expression pattern, albeit with a lower amplitude of variation for Cl3. From stage St5 onwards, the qualitative expression profile of Cl1 genes is the inverse of those of Cl2 and Cl3, suggesting that they represent opposing regulatory activities. Functional enrichment analyses revealed distinct but overlapping signatures among clusters (Figs. 3B; Supplementary Tables S9). In particular, GO terms associated with developmental processes, transcriptional regulation, and hormone-related pathways were significantly enriched, including categories related to inflorescence development, regulation of gene expression, and hormone-mediated signalling. Several transcription factor families were significantly over-represented in distinct clusters (Fig. 3C; Supplementary Tables S10), including AP2-EREBP, MADS-box, HB, ALOG, SPL, and bHLH groups, consistent with their known importance in inflorescence and meristem development. Additionally, a significant over-representation of hormone-related terms was observed (Fig. 3C). Auxin-associated categories were enriched in Cl1 and Cl2, whereas cytokinin-related terms were significantly over-represented in all three clusters, especially Cl3. Together, these results emphasize the importance of transcriptional regulation and hormone signalling in panicle differentiation and the meristem determinacy transition.

**Fig. 3:**
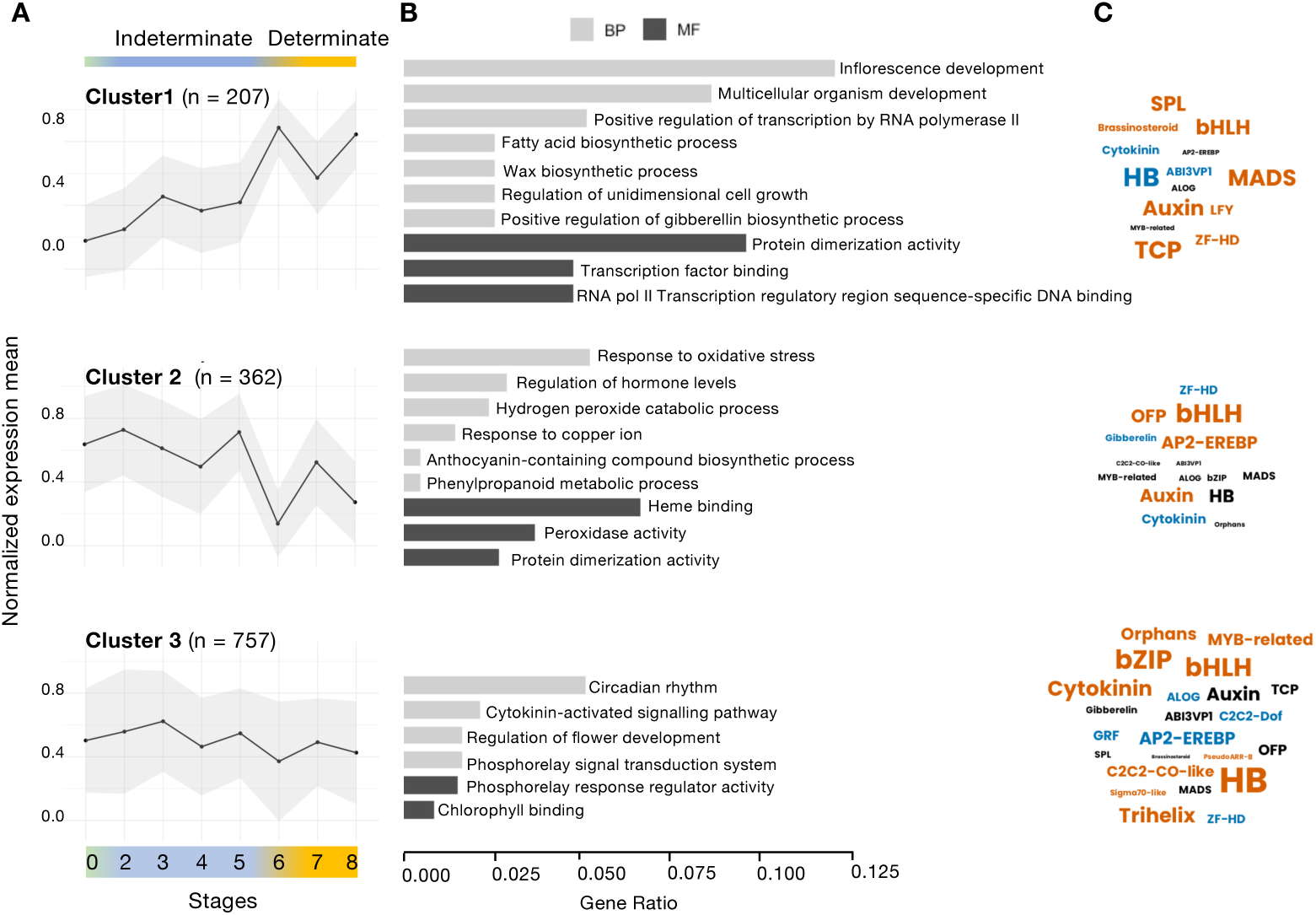
Clusters of stage-enriched genes derived from panicle differentiation associated DEGs. The transcriptional profiles corresponding to the three main co-expression networks, from clustering of 2636 DEGs, are shown. **A.** Mean gene expression profiles across developmental stages for the three clusters identified using the coseq R package. Lines represent the average normalized and scaled expression levels of genes within each cluster and surrounding ribbons indicate dispersion (standard deviation) within clusters. **B.** Selected and representative GO terms (Biological Process [BP] and Molecular Function [MF]) enriched in each cluster. The most representative and significant terms are represented (*p*.adjust <0.05) and are sorted by gene ratio. **C.** TF families and hormone-related term enrichment represented as word clouds. Word size reflects relative frequency within each cluster. Colours indicate statistical significance based on *p-*values: black, non-significant; blue, 0.01 ≤ *p-*value < 0.05; red, *p-*value < 0.01. Abbreviations (encoded protein types): ABI3VP1, ABSCISIC ACID INSENSITIVE 3 / VIVIPAROUS1; AP2-EREBP, APETALA2 / Ethylene-Responsive Element Binding Protein; bHLH, basic Helix-Loop-Helix; bZIP, basic Leucine Zipper; bZIP, basic Leucine Zipper; C2C2-CO-like, CONSTANS-like zinc finger proteins; C2C2-Dof, DNA binding with One Finger proteins; GRF, Growth-Regulating Factors; HB, Homeobox; LFY, LEAFY; MADS, MADS-box transcription factors; OFP, OVATE Family Proteins; SPL, SQUAMOSA Promoter Binding Protein-Like; TCP, TEOSINTE BRANCHED1/CYCLOIDEA/PCF family; ZF-HD, Zinc Finger–Homeodomain proteins.

### 3.3-Differential hormone accumulation during early panicle development

To obtain a complementary biochemical insight into the involvement of auxins and cytokinins in rice panicle differentiation, we compared their accumulation in whole BM- and SpM- type panicles (Fig. 4A, Supplementary Table S11). Several forms were detected for each hormone, notably IAA and its aspartate conjugate IAA-Asp in the case of auxins, as well as *cis-* and *trans-*zeatins (Zmix) in the case of cytokinins. The most abundant auxin form detected was IAA-Asp. No statistically significant differences in auxin accumulation were detected between the IM-and DM-enriched panicle states. On the other hand, a significant difference was observed between the two panicle types for *cis*- and *trans*-zeatins (Zmix), which accumulated at higher levels in the IM-enriched panicles compared to the DM-enriched type (Fig. 4A).

**Fig. 4:**
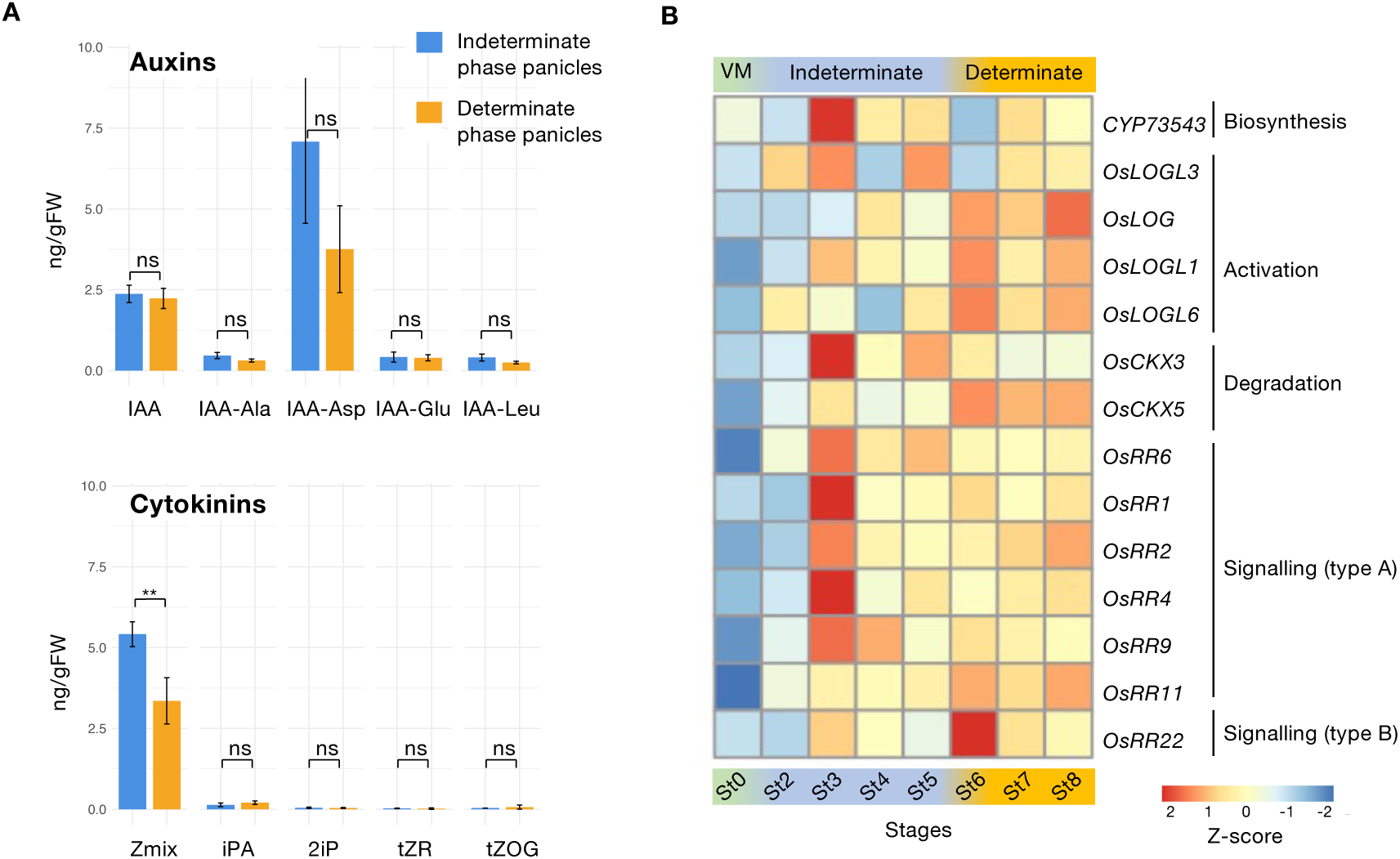
Hormonal quantification in relation to panicle development and stage specificity of cytokinin-related gene expression. **A.** Auxin and cytokinin levels in panicles at Indeterminate and Determinate phases. Data are means ± SE (n = 4; biological replicates of 4 panicles). Values are expressed in ng of hormone per g of fresh weight (ng/gFW).** *p*-values < 0.01 generated by student’s tests; Abbreviations: ns, non significant; IAA, Indole-3-Acetic Acid; IAA-Ala, N-(Indole-3-acetyl)-alanine acid; IAA-Asp, N-(Indole-3-acetyl)-aspartic acid; IAA-Glu, N-(Indole-3-acetyl)-glutamic acid; IAA-Leu, N-(Indole-3-acetyl)-leucine; Z-Mix; *cis-* and *trans*-zeatin; iPA, Isopentenyladenosine; 2iP, Isopentenyladenine; tZR, *trans*-zeatin Riboside; tZOG, *trans-*zeatin-O-Glucoside. **B.** Developmental expression profiles of cytokinin-related genes identified in the gene clustering analysis. Expression values are represented on a Z-score -2 to +2 scale. Abbreviations (activities of encoded proteins): A, activation; B, biosynthesis; D, degradation; S, signalling.

To obtain information on the regulation of cytokinin synthesis and signalling in relation to panicle branching, we next examined the expression patterns of the 14 genes annotated as cytokinin-related in the three identified gene expression clusters. Expression patterns, which were visualised as heatmaps (Fig. 4B), revealed a number of key features. Firstly, nearly all the 14 identified genes display only a basal expression in either the vegetative meristem or the St2 panicle, their activities increasing and peaking during the subsequent developmental stages. This is suggestive of a central role for cytokinins in the processes of panicle branch initiation and differentiation. Secondly, the strong peak of expression of *CYP73543* at St3 is indicative of an activation of cytokinin biosynthesis, the latter gene encoding a cytochrome P450 monooxygenase that catalyzes hydroxylation of iP-ribotides to tZ-ribotides (Takei et al., 2004). A concomitant expression increase was observed for the *LOGL3* and *LOGL1* genes encoding cytokinin-activating enzymes (Chen et al., 2022; Tokunaga et al., 2012; Zhao et al., 2024). Thirdly, the possibility of a negative feedback mechanism is suggested from the fact that the *OsCKX3* gene, encoding a cytokinin-degrading enzyme (Ashikari, 2005), displays a similarly strong increase in expression at St3. Finally, as regards the cytokinin signalling response (Brenner et al., 2012), its activation during the indeterminate panicle phase is apparent from the fact that five of the 6 type-A response regulators studied display their highest expression at St3. In contrast, the *OsRR22* gene, encoding a B-type response regulator that inhibits cytokinin signalling, displays an expression peak at the onset of panicle meristem determinacy at St6. Overall, the data in Fig. 4B highlight the importance of cytokinin biosynthesis, turnover, and associated signalling mechanisms during the key developmental stages preceding and following the panicle meristem determinacy transition.

### 3.4-Single meristem gene expression signatures for the indeterminate and determinate meristem states

Single meristem IM and DM samples obtained by panicle dissection (Figs. 5A and supplementary Fig. S5) yielded nanogram quantities of RNA for Illumina sequencing and expression analysis (Supplementary Table S12 and S13). After normalisation, 22007 genes were retained on the basis of their transcripts being detected in two or more samples from at least one of the two developmental stages (Supplementary Table S14). In total, 723 DEGs (|log2FC| > 1.0; FDR cutoff 0.05) were identified, of which 490 were IM-upregulated and 233 DM-upregulated (Fig. 5B, Supplementary Table S15). Landmark regulators of rice panicle architecture and meristem fate were identified among the DEG sets (Supplementary Table S15). In the volcano plot (Fig. 5B), 16 of these genes are highlighted, and their expression patterns confirm that the two meristem types were represented as expected.

**Fig. 5:**
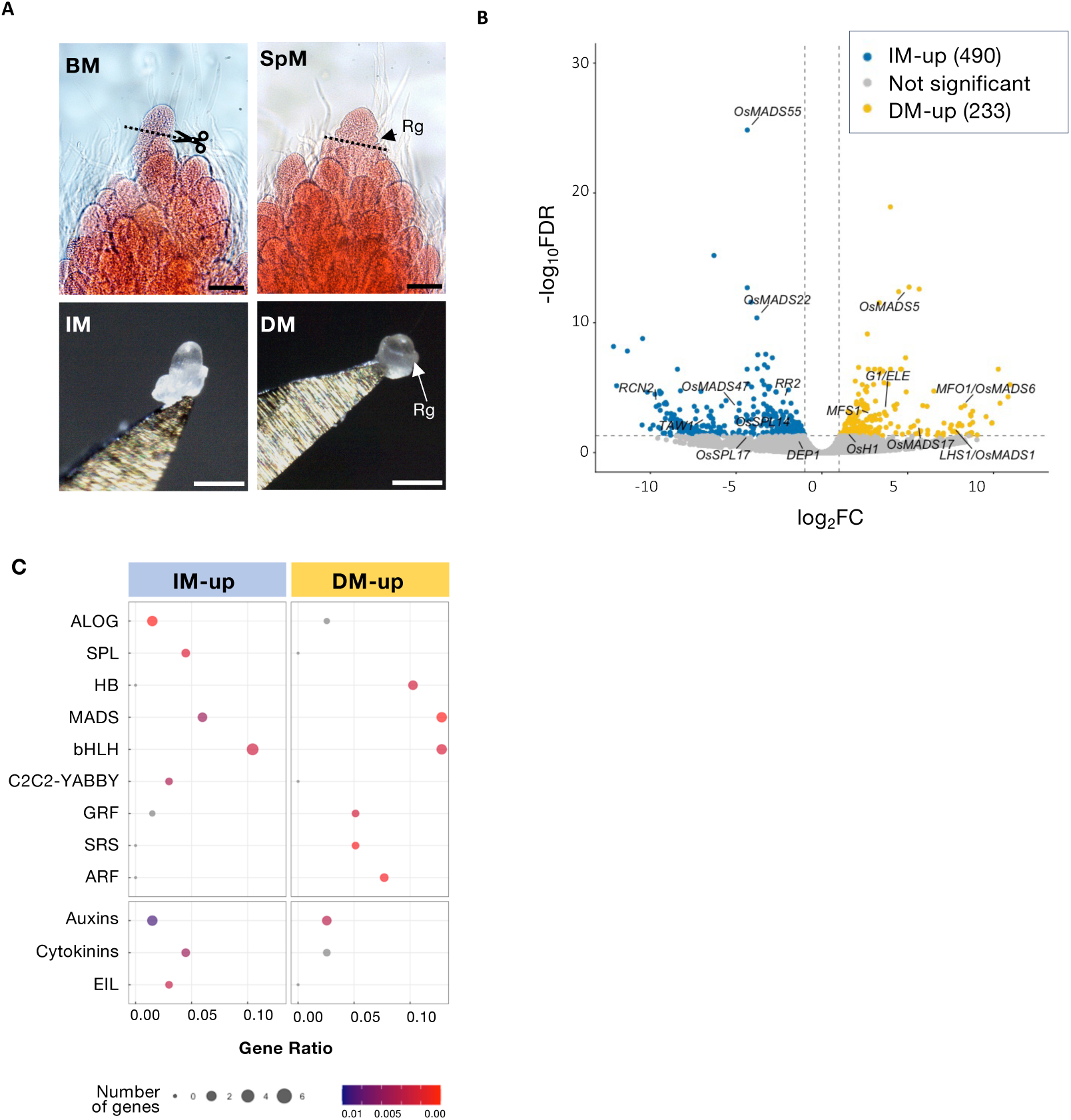
Transcriptomic analysis of individual indeterminate (IM) vs Determinate (DM) meristems of rice panicles. **A.** Images of individual Indeterminate (IM) and Determinate (DM) terminal meristems sampled from the uppermost primary branch tip of a panicle during either the branching (BM) or the spikelet meristem (SpM) phase (Rg, Rudimentary glumes). Scale bar is 100 µm. **B.** Volcano plot of differentially expressed genes (DEGs) identified between IM and DM. Each point represents the average accumulation level of transcripts across three biological replicates. Differential expression was considered significant when respectively a |log_2_ fold change| ≥ 1 (indicated by light grey dashed vertical lines) and a false discovery rate (FDR) ≤ 0.05 [corresponding to |–log10(FDR)| ≥1.3] (indicated by light grey dashed horizontal lines) were observed. Points are coloured based on mean expression across all samples as follows: blue for genes down-regulated in DM; red for genes up-regulated in DM; and grey for non-significant genes. Labelled points correspond to selected landmark genes previously described in the literature for their involvement in panicle differentiation. **C.** Analysis of DEG enrichment in transcription factor-encoding and hormone-related gene families. Plot of over-represented categories within down- and up-regulated gene sets between IM and DM samples. The most representative and significant terms are represented and sorted by gene ratio. Spot sizes indicate the number of DEGs associated with the TF and hormone-related categories; spot colour indicates the significance of the enrichment (*p*.value<0.01). Abbreviations (encoded protein types): ARF, Auxin Response Factors; bHLH, basic Helix-Loop-Helix; C2C2-YABBY, YABBY transcription factors (C2C2 zinc finger type); EIL, Ethylene-Insensitive3-LIKE; GRF, Growth-Regulating Factor; HB, Homeobox; MADS, MADS-box; SPL, SQUAMOSA Promoter-Binding Protein-Like; SRS, SHI Related Sequence.

Global gene expression trends associated with the panicle meristem determinacy transition were assessed as before by investigating GO term enrichment in DEG sets (Figures S6; Supplementary Table S16). Within the BP category, a significant enrichment (*p*.adjust < 0.05) was observed in the DEG sets of genes linked with GO terms related to "inflorescence development" (25 genes), "hormone-mediated signalling pathways" (23 genes), and "positive regulation of transcription by RNA polymerase II" (10 genes). In terms of MFs, the most significantly enriched GO terms were "DNA-binding transcription factor activity" and "sequence-specific DNA binding", corresponding to 51 and 29 genes, respectively (Supplementary Fig. S6, Supplementary Table S16). These enrichments highlight the importance of transcriptional regulation and hormone signalling in the determinacy transition. To further resolve the observed patterns, the IM- and DM-upregulated DEG sets were analysed separately. We found an enrichment of *ALOG* and *SPL* family member genes in the IM-upregulated DEG set, of HB- and ARF factor-encoding genes in the DM-upregulated group, and of genes encoding MADS-box, bHLH, and Growth-Regulating Factor (GRF) transcription factors in both (Fig. 5C, Supplementary Table S17). Given their established roles in inflorescence development, the expression patterns of the *SPL*, *ALOG*, and MADS-box family member genes were examined in greater detail (Supplementary Fig. S7). Of the 19 *SPL* genes annotated in the *O. sativa* cv. Nipponbare genome, 17 were expressed in our dataset, including four that were differentially expressed between IM and DM states, all of which were upregulated in the IM state. These included, as mentioned above, *OsSPL14*, a key regulator of panicle architecture (Jiao et al. 2010; Miura et al. 2010). Five of the *ALOG* DEGs were found to be upregulated in the IM state, including *TAW1* (Yoshida et al. 2013); while one gene, encoding the ALOG TF OsG1, is upregulated in the DM. Finally, the *O. sativa* MADS box family contains 65 genes, of which 22 (61%) were found to be expressed in our dataset, including 9 DEGs (25%). Four of these genes are upregulated in the IM state and five in the DM one.

Our analysis also revealed a notable enrichment of hormone-related genes among the DEGs, distinguishing the two developmental states (Fig. 5C, Supplementary Table S16 and S17). The expression profiles of individual DEGs associated with auxin and cytokinin pathways revealed that 5 auxin-related genes were upregulated in the IM state, 3 being implicated in signalling and 2 in biosynthesis (Supplementary Fig. S7, Supplementary Table S15). Additionally, 4 auxin-related genes showed higher expression in the DM state; in this case, they were associated with auxin signalling (1 gene), and auxin response factors (ARF) (3 genes) (Supplementary Fig. S7, Supplementary Table S15). Our analyses also revealed that the cytokinin-related DEG subset contained genes that were upregulated in both the IM and the DM states, with assigned functions in both signalling (3 IM-upregulated genes, 1 DM-upregulated gene) and in hormone activation (1 DM-upregulated gene) (Supplementary Fig. S7, Supplementary Table S15).

### 3.4-Gene regulatory network inference in relation to the IM–DM transition

Based on the preceding analyses, we constructed a gene regulatory network (GRN) to identify key regulators, at the transcriptional level, underlying the transition from an indeterminate to a determinate meristem phase during panicle differentiation. GRN inference was performed using a regression tree–based random forest approach on transcriptomic data from St2 to St7 stages, by combining DEGs from the previously defined datasets. Transcription factor-encoding and hormone-related genes were considered as candidate regulators. Putative regulatory relationships were inferred using the 437 DEGs identified in the St4–St6 and St5–St6 comparisons in the single whole panicle dataset, as well as DEGs distinguishing IM and DM states in the single-meristem dataset (|log2FC| > 2, q-value < 0.01) (Supplementary Table S18). The combined gene set was restricted to contain only genes belonging to co-expression clusters identified across panicle developmental stages. The resulting network comprises 336 nodes and 690 edges, organized into 11 communities (Supplementary Fig. S8; Supplementary Tables S19 and S20). It displays a sparse and heterogeneous architecture typical of biological GRNs, with an average degree of 4.13 and a network density of 0.012. The degree distribution shows a sharp peak at degree 1, a gradual decay up to degree 5, and a moderate tail with 31 nodes at degrees 10–20 (Supplementary Fig. S9). The many low-degree nodes and a small number of highly connected hubs collectively suggest a hierarchical organisation with a small set of highly connected regulators that structure the network into modules. Community sizes range from 2 to 50 nodes, with 1 to 118 total edges (in- and out-edges) per community (Supplementary Fig. S9). These communities remain interconnected through shared nodes and cross-community edges, indicating coordinated regulatory interactions rather than isolated functional units. Of the 336 nodes, 65 encode TFs, and 16 are hormone-related genes, forming a regulatory core potentially coordinating the IM–DM transition. In total, 44 TFs and 7 cytokinin-related genes have not been previously associated with panicle branching (Supplementary Table S20). Nevertheless, several belong to TF families, such as ALOG, AP2-EREBP, HB, and SPL, which are known to play roles in panicle architecture regulation.

Examination of node degree and neighbourhood connectivity within communities (Supplementary Fig. S9) identified several communities as being the most connected, notably communities 1, 6 and 8 which each contain clusters of regulators with components of likely significance to the IM–DM transition (Fig. 6). Community 1 predominantly brings together genes downregulated in the DM state, including five AP2-EREBP TFs—among them *IDS1* and *SNB* - previously described in relation to panicle development (Lee and An, 2012). Hormone-related genes such as *Gn1a/OsCKX2*, *OsLOGL3,* and *LAZY1* (Zhang et al., 2018) are also embedded in this module, together with two *ALOG* transcription factor genes, *OsG1L2* and *OsG1L4*, forming a tightly connected regulatory cluster. In contrast, the genes belonging to community 8 are primarily upregulated in the DM state. This module features MADS-box TF gene hubs (*OsMADS5*, *OsMADS14,* and *OsMADS34*) and cytokinin-related genes (*OsRR5*, *OsRR1,* and *OsCKX5*). Interestingly, the *ALOG* gene *OsG1L3* - which displays an opposing expression pattern (upregulated in the IM) - is connected here, suggesting cross-regulatory dynamics. The presence in the same gene community of the *APO2* and *FZP* genes that are both key determinants of inflorescence structure (Ikeda-Kawakatsu et al., 2012; Komatsu et al., 2003) suggests that interactions within this group are important to the regulation of panicle meristem activities. Moreover, it is known that the rice *SEPALLATA*-related genes *OsMADS5* and *OsMADS34*, both of which form nodes in community 8, act in cooperation to limit inflorescence branching by repressing the TERMINAL FLOWER1-Like *RCN4* (Zhu et al., 2022). The *ALOG* TF gene family is widely represented in the GRN, with members distributed across several communities (Supplementary Fig. S10). In addition to *OsG1L2* plus *OsG1L4* in community 1 and *OsG1L3* in community 8, the previously characterised *ALOG* genes *TAW1* and *OsG1* are located in community 5, while less characterised members such as *OsG1L9, OsG1L6,* and *OsG1L1* fall within communities 2, 6, and 10, respectively. This widespread integration of *ALOG* genes within distinct modules suggests that they act at multiple sites of the regulatory hierarchy controlling the IM–DM transition. Other TF families also emerge as central nodes. Members of the *OVATE* gene family are among the most connected regulators in communities 2 and 10 (Supplementary Fig. S10). For example, *OsOFP* (Lu et al., 2025) shares a community with *OsSPL8* and *OsMADS47*, placing these OVATE, SPL, and MADS-box TFs within a shared regulatory context. As regards hormone-related genes, different cytokinin and auxin pathway components are interspersed throughout several communities, further illustrating the integral role of hormonal signalling within the GRN. Notably, community 6 (Fig. 6; Supplementary Fig. S10), which shows a stronger representation of IM-upregulated DEGs, contains several hormone-related genes (*OsRR1*, *OsRR2*, *CYP735A3*, *OsLOGL9*) together with known architectural regulators such as *RCN4* and *DEP1*, suggesting that this module may contribute to maintaining branching potential during early developmental stages (Supplementary Fig. S10).

**Fig. 6:**
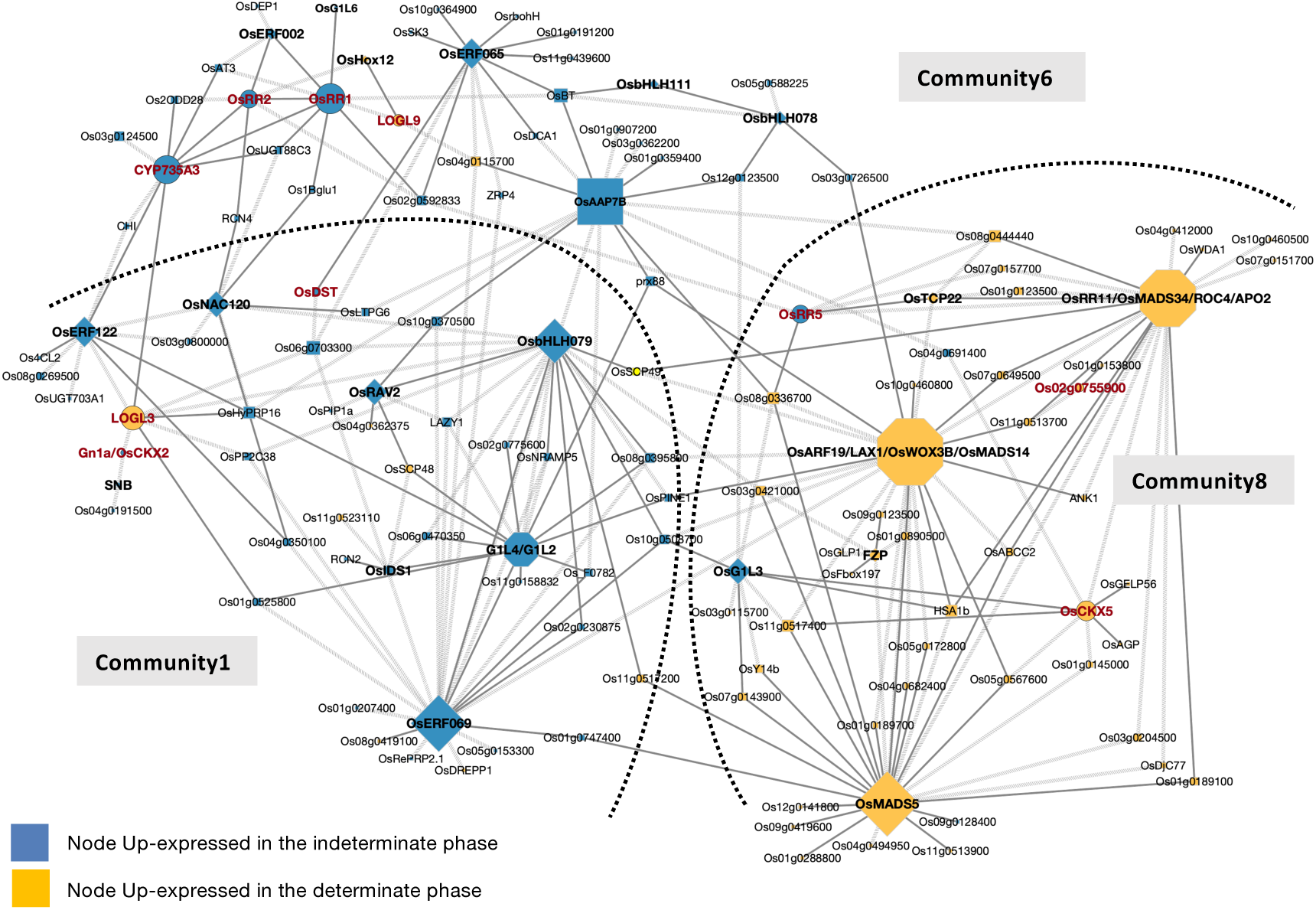
Graphical representation of sub-gene regulatory network communities 1, 6, and 8 from the inferred gene regulatory network (GRN), highlighting regulators involved in the indeterminate to determinate phase transition. Node colours indicate expression status in the determinate phase: upregulated (yellow) and downregulated (blue). Transcription factors are represented as diamonds, hormone-related genes as ellipses, groups of regulators as hexagons and other genes as rectangles. Transcription factors are shown in bold, while genes associated with the cytokinin pathway are highlighted in red and bold. Node size is proportional to degree. Solid edges correspond to weights < 0.03, whereas dashed edges (backward slash pattern) indicate weights between 0.03 and 0.05.

## 4 Discussion

### 4.1- Developmental dynamics and spatial high resolution of panicle gene expression provide a framework to describe meristem determinacy and to identify new actors

The formation of panicle branches and regulation of their developmental trajectory through meristem determinacy are processes that govern the number of spikelets per panicle and therefore yield potential. In an earlier study, we used a combined laser microdissection (LMD) and RNA-sequencing approach to compare the gene expression profiles of the RM, PbM, elongating primary branch meristem (ePbM), and SpM regions of the developing rice panicle (Harrop et al., 2016). More recently, a single-cell RNA-seq approach involving protoplasting was described by Zong et al. (2022). Both datasets provided insights into the molecular basis of rice inflorescence development, notwithstanding certain limitations in each case. With the LMD-based approach, only a few genes showing contrasting differential expression between the ePbM and SpM stages were revealed. Moreover, this analysis did not reveal any clusters of genes associated with an abrupt expression switch from the IM to DM states, although gradual decreases or increases were observed. In the case of the single-cell analysis, protoplasts were isolated from inflorescence tissues collected at various developmental stages, including both the branching and spikelet formation phases, and were compared to those from the floret stage. Although this approach may facilitate the deciphering of developmental trajectories, given that each protoplast analysed represents an individual cell, it inevitably produces expression profiles representing a continuously variable range of developmental stages. Additionally, the stress induced during protoplast isolation can lead to further variability in gene expression, potentially bringing additional complexity when developmental asynchrony and stress-related transcriptional responses interact. In this study, our two transcriptomic datasets were obtained and analysed with the aim of exploring separately and conjointly the temporal trajectory of panicle development and the discrete transition in meristem fate. The single-panicle time-course analysis identified distinct stage-dependent DEGs that govern branch development and co-expressed modules associated with successive developmental phases, whereas the single-meristem dataset was focused on the IM–DM transition at the apex of the most distal primary branch, these two approaches shedding light from complementary angles on the role of transcriptional regulation in panicle differentiation. The present study, whilst validating the importance of a number of known landmark actors, has also allowed the identification of a range of novel genes of interest that belong to families of developmental importance, but which have not to date been assigned a role in the regulation of flowering. Of particular interest are the DEGs encoding TF proteins of the MADS-box, SPL, ALOG, AP2-EREBP, and HB families (Supplementary Tables S5 and S15).

### 4.2. Dynamic transcriptional modules and transcription factor hubs

The global expression patterns observed across the panicle time-course reveal a series of stage-specific transcriptional reprogramming events, the most pronounced shifts in terms of DEG numbers occurring at the vegetative-to-reproductive transition (St0–St2) and at the indeterminate-to-determinate transition (St5–St6). Co-expression analysis further resolved these changes into three robust gene clusters, each comprising a distinct set of genes with characteristic temporal profiles, indicating that the panicle developmental program is shaped by a limited set of transcriptional modules, acting with stage-specificity to govern meristem states.

One of the most striking findings is the consistent enrichment of TF gene families across both datasets, within the co-expression clusters of the panicle time-course and among the DEGs distinguishing meristem types. This suggests that local shifts in meristem identity guide panicle morphogenesis through the action of transcriptional regulatory backbones. Cluster 1, which shows an overall rising trend in expression and peaks at St6 and St8, is particularly enriched in genes encoding reproductive regulator proteins, including multiple MADS-box genes with well-characterized roles in rice inflorescence development: *LHS1*/*OsMADS1* controls floret meristem specification and regulates palea and lemma differentiation (Khanday et al., 2013; Chen et al., 2006). *OsMADS5* and *OsMADS34* function cooperatively to limit inflorescence branching (Zhu et al., 2022), while *MOSAIC FLORAL ORGANS1 (MFO1)*/*OsMADS6* regulates floral organ identity and meristem fate (Ohmori et al., 2009). OsMADS14 and *OsMADS15* participate in inflorescence meristem identity specification and vegetative branching (Lu et al., 2012; Kobayashi et al., 2012). Additional Cluster 1 members known to be determinants of inflorescence structure include *LAX PANICLE1* (*OsLAX1*), encoding a bHLH protein that regulates rachis-branch and spikelet development (Komatsu et al., 2001), and *OsSPL7*, which participates in regulating both panicle architecture and tiller number (Yan et al., 2021). Their collective upregulation suggests that Cluster 1 acts as a coordinated module that concomitantly promotes floral identity and terminates branching.

Clusters 2 and 3 reveal contrasting TF gene enrichments in relation to the branching stages of development. Cluster 2 shows AP2-EREBP family overrepresentation, while Cluster 3 is enriched in HB family factors, mirroring patterns described by Harrop et al. (2016) during early branch meristem activity. This convergence between our clustering and previous LMD data supports the importance of these transcriptional modules in sustaining indeterminate growth. By complementing the temporal profiles, our tissue-specific single meristem dataset allows the identification of a number of additional TF gene family members as potential regulators of meristem determinacy (Supplementary Table S15). All five *SPL* DEGs were observed to be more expressed in the IM state, including the extensively described panicle development gene *OsSPL14* (Jiao et al., 2010; Miura et al., 2010). The other four SPL-encoding DEGs (*OsSPL17*, *OsSPL7*, *OsSPL10* and *OsSPL3*) have been mostly described in terms of other developmental functions: *OsSPL3* (a regulator of *OsMADS50*) in crown root development (Shao et al., 2019); *OsSPL7* in tillering (Dai et al 2018; Wang and Zhang 2017; Yan et al., 2021,); *OsSPL10* in trichome formation (Lan et al., 2019 G3). The similar upregulation of expression observed for all 5 genes with respect to meristem indeterminacy suggests previously undescribed roles for some members of the group in the promotion of inflorescence branching. Our results also reinforce the central role of *ALOG* genes in the regulation of rice panicle development and suggest that their functions include the transition phase of meristem fate, consistent with previous studies on the *ALOG* family members *TAW1*, *OsG1L1,* and *OsG1L2* (Yoshida et al. 2013; Beretta et al. 2023; Luo et al. 2025). The observed differential expression in our datasets of four additional *ALOG* genes (*OsG1L3*, *OsG1L4*, *OsG1L6,* and *OsG1L9*) might be an indication of either an overlapping role with *TAW1* and/or a potential functional diversification within the *ALOG* family. The fact that these *ALOG* members show different expression clustering profiles (Cl1 for *OsG1L4*, Cl2 for *OsG1L3*, unclustered for *TAW1*, *OsG1L6,* and *OsG1L9*) suggests a diversification at least in terms of temporal activities. More generally, GRN analysis reinforces the abovementioned findings by positioning multiple TFs as hub nodes within IM-or DM-associated communities (Fig. 6). Notably, the IM-associated community 1 integrates AP2-EREBP TF genes (*IDS1, SNB, OsERF069, OsERF12221, OsRAV2*), and ALOG (*OsG1L2, OsG1L4*) regulators along with hormone-related genes (*Gn1a/OsCKX2*, *OsLOGL3*), while DM-associated community 8 centres on MADS-box hubs (*OsMADS5*, *OsMADS14*, *OsMADS34*) interacting with *OsG1L3* and cytokinin regulators (*OsRR5*, *OsRR1*). Members of the OVATE family, such as *OsOFP,* which is clustered with *OsSPL8* and *OsMADS47* in other communities, emerge as connectors linking TF and hormone modules. Together, these network properties suggest that TFs act as integrative hubs that coordinate developmental timing, meristem identity, and hormonal inputs during the IM–DM transition.

### 4.3-A central role for auxin and cytokinin in rice panicle development

Among the complex interactions that operate between hormones and regulatory factors, the balance between auxin and cytokinin, at local and regional levels, is a key parameter that influences the development of the plant (Schaller et al., 2015). Our combined biochemical and transcriptomic analyses indicate that rice panicle development, and in particular the transition from IM to DM state, is accompanied by a progressive hormonal reprogramming.

#### Gene expression profiling reinforces the key role for auxin in panicle differentiation

Auxin accumulation, at the level of the panicle, did not show a statistically significant difference between the panicle types (BM phase vs. SpM phase). However, transcriptomic analyses revealed extensive differential regulation of auxin-related genes across developmental transitions, including the vegetative-to-reproductive phase and the shift from IM to DM state. Auxin-related terms were enriched both in stage-specific DEG sets and in co-expression clusters associated with a sharp expression change around the indeterminate to determinate phase transition (i.e., Cl1 and Cl2), indicating overall that auxin signalling operates throughout panicle development. Several genes associated with auxin transport and signalling are upregulated at the onset of determinate development (St6) in the single-panicle dataset; namely, *OsIAA26*, *OsPIN1C,* and *OsPIN1D*. The single-meristem dataset provided complementary tissue-specific information on the IM-DM transition, revealing that several genes associated with auxin biosynthesis and primary response show preferential expression in indeterminate meristems, including *OsYUCCA10*, a cytochrome P450 likely involved in auxin production (Zong et al., 2022), and *OsSAUR8*, suggesting enhanced auxin responsiveness in proliferative meristems. Multiple gene regulators of auxin signal transduction were also differentially expressed between IM and DM states, including three *AUXIN RESPONSE FACTOR* genes (*OsARF2*, *OsARF7a*, *OsARF15*) predicted to encode transcriptional repressors (Song et al., 2023), and four AUX/IAA genes (*OsIAA2*, *OsIAA24*, *OsIAA30*, *OsIAA31*) displaying contrasting temporal expression patterns suggestive of functional diversification. ARFs bind to auxin response elements (AuxREs) found in the promoters of auxin-responsive genes to exert either transcriptional activation or repression of expression (Liu et al., 1994). Aux/IAA proteins inhibit promoter-bound ARFs by recruiting corepressor complexes (Ulmasov et al., 1997). Because these factors modulate the transcriptional output of auxin pathways, such shifts likely alter auxin sensitivity and downstream gene regulation in the absence of drastic changes in auxin abundance. Moreover, the highly localized nature of auxin gradients observed in meristematic tissues (Yang et al., 2017; Liu et al., 2021; Sato et al., 2025) has demonstrated that spatial distribution and transport dynamics can be more informative than bulk hormone concentration. The importance of local auxin gradients in the regulation of inflorescence differentiation is illustrated by the *PLANT ARCHITECTURE AND YIELD1* (*PAY1)* gene involved in polar auxin transport, mutants of which display an altered panicle phenotype (Zhao et al.,2015).

#### Cytokinin dynamics support indeterminate meristem activity during panicle branching

In contrast to the situation observed for auxin, a significantly higher level of cytokinin (zeatin) was measured in developing panicles during the BM phase (enriched in IM-type meristems) compared to the SpM phase (enriched in DM-type meristems). This observation provides a global view of hormone accumulation and aligns well with the known role of cytokinins in plants, namely in maintaining stem cell activity, meristem size, and meristem identity, thereby affecting inflorescence branching (Kyozuka, 2007; Zhao, 2008). Transcriptomic data further support this interpretation by revealing a coordinated regulation of genes responsible for cytokinin biosynthesis, degradation, and signalling across developmental stages and with respect to meristem identities. Genes encoding enzymes involved in cytokinin production were preferentially expressed during active branching phases. In particular, the *CYP735A3* gene, responsible for the production of *trans*-zeatin–type cytokinins, displayed a sharp expression peak at stage St3, thus coinciding with intense branch meristem formation seen at the morphological level and with elevated zeatin levels detected during the indeterminate phase of development. The promotion of cytokinin biosynthesis is also suggested by the presence within the co-expression clusters of several members of the *LONELY GUY-like* (*LOGL*) family encoding enzymes that convert cytokinin ribotides into bioactive free bases, namely *OsLOG*, *OsLOGL1*, *OsLOGL3,* and *OsLOGL6*. Our RNA-seq data revealed a diversity of expression profiles within this family, a majority of which (6/11 genes, including *OsLOGL3*) were more highly expressed during indeterminate stages St3-St5. Moreover, cytokinin turnover appears to be tightly regulated, as witnessed by the presence of two cytokinin oxidase/dehydrogenase genes, *OsCKX3* and *OsCKX5*, in the expression clusters. The sharp expression peak of *OsCKX3* at stage St3 mirrors that of *CYP735A3*, suggesting feedback control between cytokinin synthesis and degradation. In contrast, *OsCKX5* displayed a more stable expression pattern, consistent with a broader role in cytokinin homeostasis throughout development. From analyses performed on wheat, Chen et al. (2022) pointed out the similar sequence conservation of cytokinin regulatory elements (CREs) in the promoters of *LOGL* and *CKX* genes, probably reflecting the key role of CKXs in maintaining cytokinin homeostasis. The coordinated expression of *LOGL* and *CKX* genes supports the existence of a dynamic mechanism controlling cytokinin availability during branching.

The expression patterns of genes encoding cytokinin response components provide further illustration of the importance of this hormone in developmental tuning. Several type-A response regulator genes (*OsRR1*, *OsRR2*, *OsRR4*, *OsRR5*, *OsRR6*, *OsRR9*) exhibited stage-dependent expression peaks and were preferentially expressed in indeterminate meristems. Type-A response regulator genes, which were first described on account of their induction by cytokinin (Hwang and Sheen, 2001), encode negative regulators of the upstream cytokinin signal transduction system, thereby participating in negative feedback. Conversely, type-B response regulator genes (Sakai et al., 2001) encode MYB-like transcription factors that activate cytokinin-responsive gene expression, thereby opposing the action of the type-A genes (Brenner and Schmülling, 2015). As type-A regulators function as primary cytokinin-inducible negative feedback elements, their expression likely reflects locally elevated cytokinin activity in proliferative meristematic regions during the indeterminate phase of panicle development. In contrast, the type-B response regulator genes *OsRR22* and *OsRR26* showed higher expression in the determinate panicle phase and DM state, respectively, suggesting a shift in the cytokinin signalling balance as meristems progress towards determinacy. Interestingly, a recent study has shown that type-A response regulators, particularly *OsRR2* and *OsRR4*, positively influence panicle development by promoting secondary branch and spikelet formation. They achieve this by modulating multiple hormonal pathways, hormone contents, and processes related to sugar metabolism, nutrient transport, cell division, and reactive oxygen species (ROS) homeostasis (Rong et al.,2025).

#### GRNs associate hormonal and other TFs to describe panicle determinacy

The key GRN communities described in the present study integrate hormonal regulators with TF hubs, suggesting that hormone signalling is embedded within the broader regulatory network that controls the IM–DM transition. Interestingly, cytokinin-related genes are distributed across several GRN communities, notably communities 1, 6, and 8, suggesting multiple regulatory contexts for cytokinin signalling during the IM–DM transition rather than a single linear pathway. Community 1 (IM-associated) connects cytokinin-related genes such as *Gn1a/OsCKX2* and *OsLOGL3* with *AP2-EREBP*, *SPL,* and *ALOG* (*G1L2*/*G1L4*) TF gene families, while community 8 (DM-associated) links *OsRR5, OsRR1*, *OsCKX5,* and *OsARF18* with MADS-box hubs including *OsMADS5*, *OsMADS14,* and *OsMADS34*. Community 5, which is largely composed of genes down-regulated during the indeterminate stage, includes *OsG1L6*, *OsERF065*, *RCN4,* and *DEP1*. This community contains a high proportion of genes that remain poorly characterised, highlighting its interest as a possible source of novel candidate regulators of meristem determinacy. The observation of GRN communities integrating TF hubs with hormonal regulators is consistent with reports of conserved dynamic transcriptional modules identified across cereal inflorescences (Li et al., 2018; Zhu et al., 2018; Van Gessel et al., 2022; Eveland et al., 2018), wherein expression shifts within stage-specific clusters and dynamic cytokinin/auxin reprogramming similarly coordinate meristem fate transitions from branching to spikelet/floret specification.

## 5 Conclusions

By associating a detailed developmental time-course of panicle development with stage-resolved meristem sampling to investigate global gene expression programs, in tandem with biochemical measurements of hormone accumulation and GRN inference, we delineated dynamic transcriptional modules, hormone-related activities, and regulatory hubs associated with the indeterminate-to-determinate transition in rice panicles. Our data not only recover the expected behaviour of known inflorescence regulators, but also identify additional members of the *SPL*, *ALOG*, and MADS-box families, amongst others, as candidate hubs within IM- and DM-associated communities (Fig. 7). In total, 44 transcription factor genes and 7 cytokinin-related genes that to date have not been functionally described in the context of rice flowering, were identified as displaying expression profiles linked to the key stages of panicle differentiation (Supplementary Table S20) and are thus of particular interest for future studies. Our combined biochemical and transcriptomic evidence supports a model in which cytokinin signalling is most active during the proliferative branching phase, during which it promotes meristem maintenance while simultaneously triggering feedback mechanisms, with progressive rewiring of auxin signalling pathways also likely to contribute to spatial organization and to the acquisition of the determinate meristem fate (Fig. 7). A better understanding of the local regulation of meristematic activities during panicle differentiation might be achieved by the use of hormone biosensor markers (Yang et al., 2017, Sato et al., 2025) to monitor the dynamic regulation of auxin and cytokinin responses in the tissues of interest. Future analyses of panicle expression dynamics in relation to genetic diversity could reveal in further detail how meristem determinacy transitions shape inflorescence differentiation and govern the consequent effects on productivity.

**Fig. 7:**
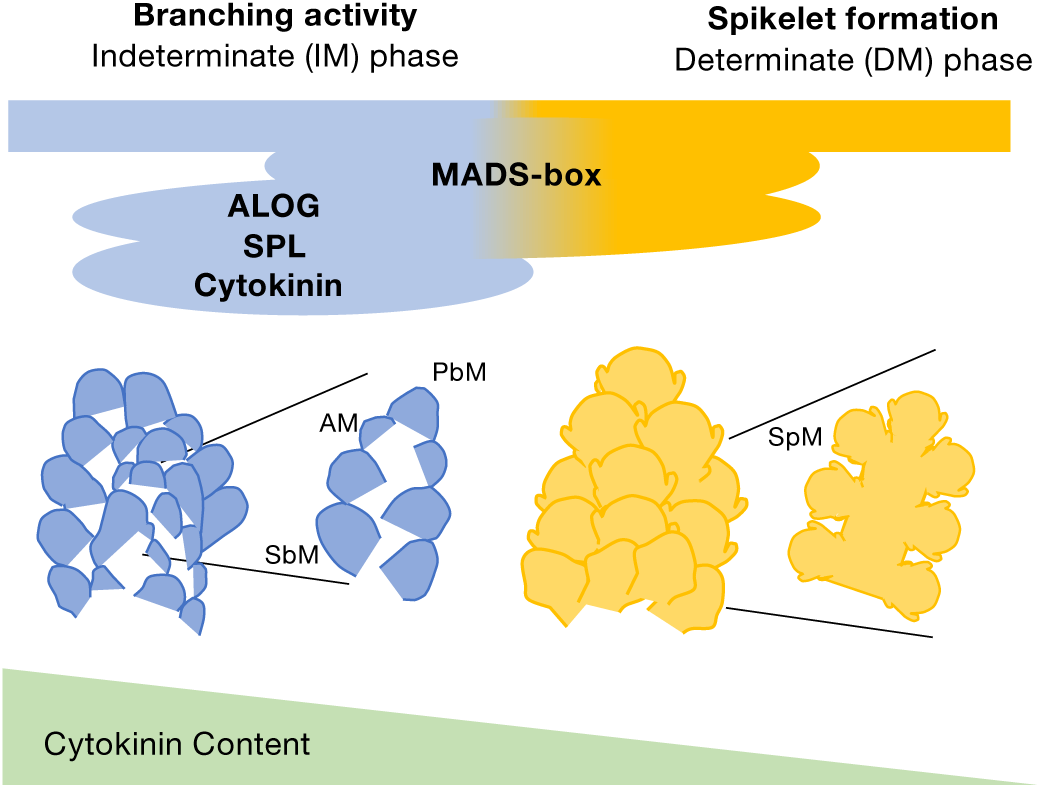
Schematic representation of key regulatory factors associated with the indeterminate to determinate phase transition during rice panicle development. Expression analyses highlighted enrichment of transcription factor-encoding and cytokinin-related genes in DEG sets associated with the progression of panicle differentiation. *MADS-box* members (*OsMADS20/55/47/22*) are up-regulated in the IM phase, while *OsMADS32/14/24/5/15* show higher expression in the DM phase. *ALOG* genes (*OsG1L1/2/3/4/6/8/9*) are preferentially expressed in the IM phase, whereas *OsG1* is up-regulated in the DM phase. *SPL* genes (*OsSPL3/5/14*) display IM-enriched expression, while *OsSPL7* expression increases in the DM phase. Cytokinin signalling type-A response regulators are upregulated in the IM phase, consistent with the higher cytokinin levels measured during this phase of panicle development, which also concord with the IM phase expression peak of the cytokinin biosynthetic gene *CYP735A3*.

## Abbreviations

AM: Axillary Meristem;
BM: Branching Meristem;
DM: Determinate Meristem;
FlM: Floret Meristem;
FlO: Floral Organ initiation;
GRN: Gene Regulatory Network;
IM: Indeterminate Meristem;
PbM: Primary Branch Meristem;
RM: Rachis Meristem;
SbM: Secondary Branch Meristem;
SpM: Spikelet Meristem;
TF: Transcription Factors;
VM: Vegetative Meristem.

## Author contributions

HA and SJ contributed to the conception and design of the experiments. HA and DA were involved in designing the RNA extraction protocol, library preparation, and sequencing strategy. JS performed the library preparation, while CG and MCC collected the plant panicles and meristems. CM and CG carried out the RNA extractions, and VV conducted the hormone analyses. JT and FB performed the RNA-seq data processing and mapping. HA analysed the data, HA and JT interpreted the results, and wrote the manuscript. SJ reviewed and edited the manuscript. All authors read and approved the final version of the manuscript.

## Acknowledgements

The authors acknowledge the ISO 9001 certified IRD i-Trop HPC (member of the South Green Platform) at IRD Montpellier for providing HPC resources that have contributed to the research results reported within this paper (URL:http://www.southgreen.fr), Alexandre Soriano (UMR AGAP, Montpellier) for his helpful advice on expression data analysis, and Pierre Serin for technical assistance in the greenhouse.

## Data availability and FAIR (Findable Accessible Interoperable Reusable) compliance statement

All data presented in this research are available in the article and supporting information. The complete read count data have been deposited in Supplementary Tables S3 and S12. The data for this study have been deposited in the European Nucleotide Archive (ENA) at EMBL-EBI under accession number PRJEBxxxx (in progress).

## Supporting informations

**Supplementary Fig. S1:** Sampling of immature panicles at nine developmental stages in three replicates.

**Supplementary Fig. S2:** Histograms showing *p*-values for each pairwise comparison of differential gene expression in relation to panicle stage.

**Supplementary Fig. S3**: Enrichment analysis of GO terms, transcription factors, and hormone-related categories in DEG sets identified through sequential pairwise developmental stage comparisons.

**Supplementary Fig. S4:** Clustering analysis and distribution of biological profiles accross clusters and stages

**Supplementary Fig. S5:** Images of the collected indeterminate (IM) and determinate (DM) meristem samples at branch (BM) and spikelet (SpM) panicle phases respectively.

**Supplementary Fig. S6:** Plots of over-represented Go-BP (biological process) and Go-MF (molecular function) of DEGs between IM and DM stages (|log_2_ FC| >1, *p*<0.05).

**Supplementary Fig. S7**: Profiling, for the IM and DM states, of differential expression of genes encoding SPL, ALOG, and MADS-box transcription factors, plus genes associated with auxin (yellow dot) and cytokinin (violet dot) pathways.

**Supplementary Fig. S8 :** Graphic representation of the inferred Gene Regulatory Network (GRN) illustrating regulators of the indeterminate to determinate phase transition.

**Supplementary Fig. S9**: Degree distribution and modular structure of the inferred Gene Regulatory Network.

**Supplementary Fig. S10 :** Representation of each Sub-GRN Community. Node colors indicate expression status in DM phase: upregulated (yellow) and downregulated (blue).

**Supplementary Table S1**: MRM parameters for each auxin and cytokinin compound

**Supplementary Table S2:** Read and mapping statistics for all RNA-seq libraries used in this study

**Supplementary Table S3:** Raw Count values for each replicate for Single panicle samples

**Supplementary Table S4:** Normalized Counts for each replicate after count filtering for the Single Panicle dataset

**Supplementary Table S5:** Differential expression test results for each pair of successive devlopmental stageswith an FDR<0.01 and |log2FC| >1. Abbreviations: log2FC, log2 Fold Change; logCPM, log10 for CountsperMillion; and FDR, False Discovery Rate.

**Supplementary Table S6:** Significant Go-BP (Biological Process) and GO-MF (Molecular Function) enrichment Terms of the DEGs identified between pariwise comparisons (p.adjust<0.05)

**Supplementary Table S7:** Transcription factor family and hormone categories enrichment analysis (p-value<0.05) of the DEGs in pairwise comparisons.

**Supplementary Table S8:** Complete list of the genes clustered

**Supplementary Table S9:** Go-BP (Biological Process) and -MF (Molecular Function) enrichments of the genes clustered (p.adjust<0.05)

**Supplementary Table S10:** Transcription factor family and hormone categories enrichment analysis of the clustered genes

**Supplementary Table S11:** Hormone quantification in ng of hormone per g of fresh weight (ng/gFW). Abbreviations: IAA, Indole-3-Acetic Acid; IAA_Asp, N-(Indole-3-acetyl)-aspartic acid; IAA_Ala, N-(Indole-3--acetyl)-alanine acid; IAA_Glu, N-(Indole-3-acetyl)-glutamic acid; IAA_Leu, N-(Indole-3-acetyl)-leucine; Z-Mix, for *cis*- and *trans*-zeatin; iPA, Isopentenyladenosine; 2iP, isopentenyladenine; tZR, trans-zeatin riboside; tZOG, trans-zeatin-o-glucoside

**Supplementary Table S12:** Read and mapping statistics for RNA-seq libraries of Single meristem samples used in this study

**Supplementary Table S13:** Raw Count values for each replicate for single meristem samples

**Supplementary Table S14:** Normalized Counts for each replicate after count filtering for single meristem dataset

**Supplementary Table S15:** Differential expression test results between IM and DM stages of Single meristem dataset. In green, genes that are DE with an FDR<0.05 and |log2FC| >1. Abbreviations: log2FC, log2 Fold Change; logCPM, log10 for CountsperMillion; and FDR, False Discovery Rate.

**Supplementary Table S16:** Significant Go-BP (Biological Process), GO-MF (Molecular Function) and KEGG enrichment Terms of the DEGs identified between IM and DM stages (p.adjust<0.05)

**Supplementary Table S17:** Transcription factor family and hormone categories enrichment analysis of the genes Down- and Up-regulated in DM.

**Supplementary Table S18:** List of genes used to infer the Gene Regulatory Network (GRN). Genes highlighed in green are described in the litterature.

**Supplementary Table S19:** List of Edges in the Gene Regulatory Network (GRN)

**Supplementary Table S20.:** List of nodes included in the inferred Gene Regulatory Network. Genes highlighed in green are described in the litterature.

